# The important choice of reference environment in microevolutionary climate response predictions

**DOI:** 10.1101/2022.01.07.475361

**Authors:** Rolf Ergon

## Abstract

It is well documented that individuals of wild populations can adjust to climate change by means of phenotypic plasticity, but few reports on adaptation by means of genetically based microevolution caused by selection. Disentanglement of these separate effects requires that the environmental zero-point (reference environment) is defined, and this should not be done arbitrarily. The problem is that an error in the environmental zero-point may lead to large errors in predicted microevolution. Together with parameter values, the zero-point can be estimated from environmental, phenotypic and fitness data. A prediction error method for this purpose is described, with the feasibility shown by simulations. As shown in a toy example, an estimated environmental zero-point may have large errors, especially for small populations. This may still be a better choice than use of an initial environmental value in a recorded time series, or the mean value, which is often used. Another alternative may be to use the mean value of a past and stationary stochastic environment, which the population is judged to have been fully adapted to, in the sense that the expected geometric mean fitness was at a global maximum. Exceptions are cases with constant phenotypic plasticity, where the microevolutionary changes per generation follow directly from phenotypic and environmental data, independent of the chosen environmental zero-point.

## 1 Introduction

Wild populations respond to changing environments by means of phenotypic plasticity and microevolution, and especially climate change responses have been extensively studied. The aim is then to disentangle phenotypic changes owing to genetically based microevolution, caused by natural selection, and changes due to individual phenotypic plasticity. Relying on 11 review articles, including reviews of altogether 66 field studies, Merilä & Hendry (2014) arrived at the conclusion that evidence for genetic adaptation to climate change has been found in some systems, but that such evidence is relatively scarce. They also concluded that more studies were needed, and that these must employ better inferential methods. The aim of the present article is to give a contribution in that respect.

It is obvious that for all evolutionary systems with interval-scaled environmental variables *u_t_*, as for example temperature in °C, a suitable zero-point (reference environment) *u_ref_* must be chosen, and as argued in Section 2 this should not be done arbitrarily. A zero-point is in general defined as “the point on a scale that denotes zero and from which positive and negative readings can be made” (Collins English Dictionary). The problem is that an error in the environmental zero-point may lead to large errors in predicted microevolution. In most cases where the environmental variable is for example a temperature, the zero-point should not, for example, be set to 0 °C. Neither should it without further consideration be set to the initial or mean environmental value of a specific time series. It appears that the need for a proper environmental zero-point definition, and thus an environmental cue definition, has been largely ignored in the reviewed studies referred to in Merilä & Hendry (2014).

The present article is an attempt to clarify some important questions relating to environmental zero-points, and for that purpose a method for model-based predictions of microevolutionary changes is also proposed. This method is based on parameter estimation by means of prediction error minimization, including estimation of the environmental zero-point and initial mean values of quantitative traits.

For a discussion of the general microevolution vs. plasticity disentanglement problem, we may for simplicity assume the intercept-slope individual reaction norm model

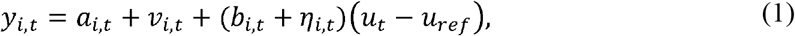

where *u_t_* – *u_ref_* and *y_i,t_* are the environmental cue and the individual phenotypic value, respectively, as functions of time *t* measured in generations. Here, the traits *a_i,t_* and *b_i,t_* are the additive genetic components of the individual reaction norm intercept and plasticity slope, respectively, while *v_i,t_* and *η_i,t_* are independent iid zero mean normal non-additive effects. The generations are assumed to be non-overlapping.

From Equation (1) follows the mean trait reaction norm model 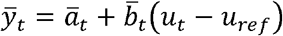, and from this simple equation follows the basic questions discussed in this article. How can the evolution of 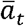 and 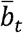 be predicted, provided that *u_t_* and *y_i,t_* are known? And how will these predictions be affected by errors in the estimated or assumed value 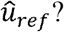 It turns out that in order to answer these questions we also need information on individual fitness values *W_i,t_*.

The environmental zero-point *u_ref_* is determined by the environment at which the phenotypic variance has its minimum, as defined in more detail in Section 2, and as discussed in Ergon & Ergon (2017) and Ergon (2018). In theoretical work it is often assumed that the population has fully adapted to a stationary stochastic environment with a given mean value, such that the expected geometric mean fitness is at a global maximum, and the environmental zero-point is then set to zero (Lande, 2009; Chevin & Lande, 2015). Although there is nothing wrong with this theoretical approach, it disguises the underlying problem discussed here, and *u_ref_* is therefore included in Equation (1). This formulation also makes it possible to distinguish between the environment as such, and the environmental cue. In some cases, it may be possible to determine the environmental zero-point experimentally, see, e.g., Fossen *et al*. (2018), but that may obviously be difficult for wild populations.

When the environmental cue *u_t_* – *u_ref_* changes over time, the mean trait values 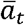 and 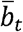 as follow from Equation (1) may evolve due to selection, and as a result also the mean phenotypic value 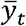 will evolve (Lande, 2009). Without changes due to selection, i.e., if the mean trait values 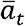 and 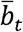 are constant, the value of 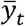 may still change when *u_t_* – *u_ref_* changes, as also follows from Equation (1).

Section 2 discusses several aspects of the general microevolution vs. plasticity disentanglement problem. First, a definition of the environmental zero-point *u_ref_* is given. Second, it is shown how the mean trait values 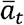 and 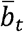, and thus also 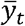, evolve as functions of the environmental cue *u_t_* – *u_ref_* and the phenotypic selection gradient *β_y,t_*. Third, it is shown how *u_ref_* and *β_y,t_*, as well as initial mean trait values and the parameter values in the ***G*** matrix, can be estimated by means of a prediction error minimization method (Ljung, 2002), using data from known time series of *u_t_* and *y_i,t_*, as well as of individual fitness values *W_i,t_.* Forth, it is discussed why it may be difficult to estimate *u_ref_*, as revealed by simulations, and which consequences errors in estimated values of *u_ref_* will have. Exceptions are here cases with constant phenotypic plasticity, where the microevolutionary changes per generation follow directly from phenotypic and environmental data, independent of the chosen environmental zero-point.

It must be underlined that the theory in Section 2 assumes that the phenotypic trait *y_i,t_* in Equation (1) is not correlated with other phenotypic traits having causal effects on fitness, see Morrissey *et al*. (2010) for a discussion. Also note that the need for a proper environmental zero-point is not specific for the simple case according to Equation (1).

Simulations in Section 3 make use of a toy example, utilizing the intercept-slope reaction norm model in Equation (1). The environmental input *u_t_* is here a noisy positive trend in spring temperature, while the individual phenotypic values *y_i,t_* are the clutch-initiation dates for a certain bird species. The toy example also assumes that the individual fitness values *W_i,t_* are the numbers of offspring. The essential questions are how microevolutionary changes in mean intercept and plasticity slope can be predicted, and how these predictions are affected by errors in the environmental zero-point *u_ref_* in Equation (1). The simulations show that errors in the estimated or assumed value of *u_ref_* may cause large mean trait prediction errors. They also show the feasibility of the proposed parameter estimation method.

Finally follows a discussion in Section 4. Derivations of prediction equations, simulation results with modeling error and increased population size, and a short comparison with REML parameter estimation are given in Appendix 1 to 4.

## 2 Theory and methods

### 2.1 Example system

For a study of the general environmental zero-point problem, and for a test of the proposed parameter estimation method, we may consider a true evolutionary system based on Equation (1),

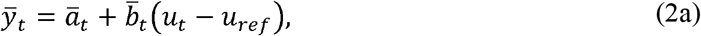

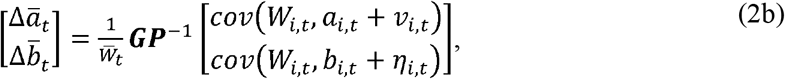

with the additive genetic covariance matrix 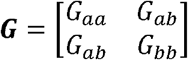, and the phenotypic covariance matrix and 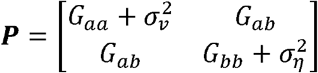. Here, Equation (2b) is the multivariate breeder’s equation (Lande, 1979), where *W_i,t_* is found from any given fitness function. It is assumed that the phenotypic trait *y_i,t_* in Equation (1) is not correlated with other phenotypic traits having causal effects on fitness, and that generations are non-overlapping.

### 2.2 Environmental zero-point

As discussed in the Introduction, there is a need for a well-defined environmental zero-point:

#### Definition 1

*Assuming a single environmental variable u_t_, and given a reaction norm model, the environmental zero-point is*

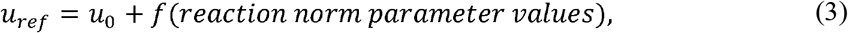

*where u*_0_ *is the environment at which the phenotypic variance is at a minimum, and where the covariance between the plastic phenotypic value and reaction norm slope is zero. Here, f (reaction norm parameter values’) is a correction term that may be zero*.

For the reaction norm model (1) we find for example (using *u*’ = *u* – *u_ref_*)

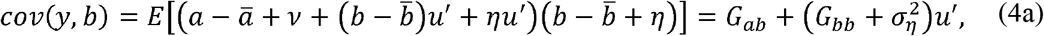

which by setting *cov*(*y, b*) = 0 and *u*’ = *u*_0_ – *u_ref_* gives the environmental zero-point

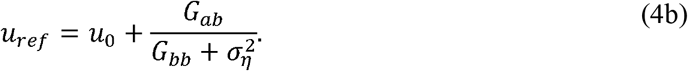

For *G_ab_* = 0 the environmental zero-point is thus the environment where the phenotypic variance is minimized (see Fig. 1 for illustration). This is also the environment where the expected geometric mean fitness has a global maximum, and thus the environment the population is fully adapted to. In this environment the environmental cue will be zero.

**Fig. 1.**
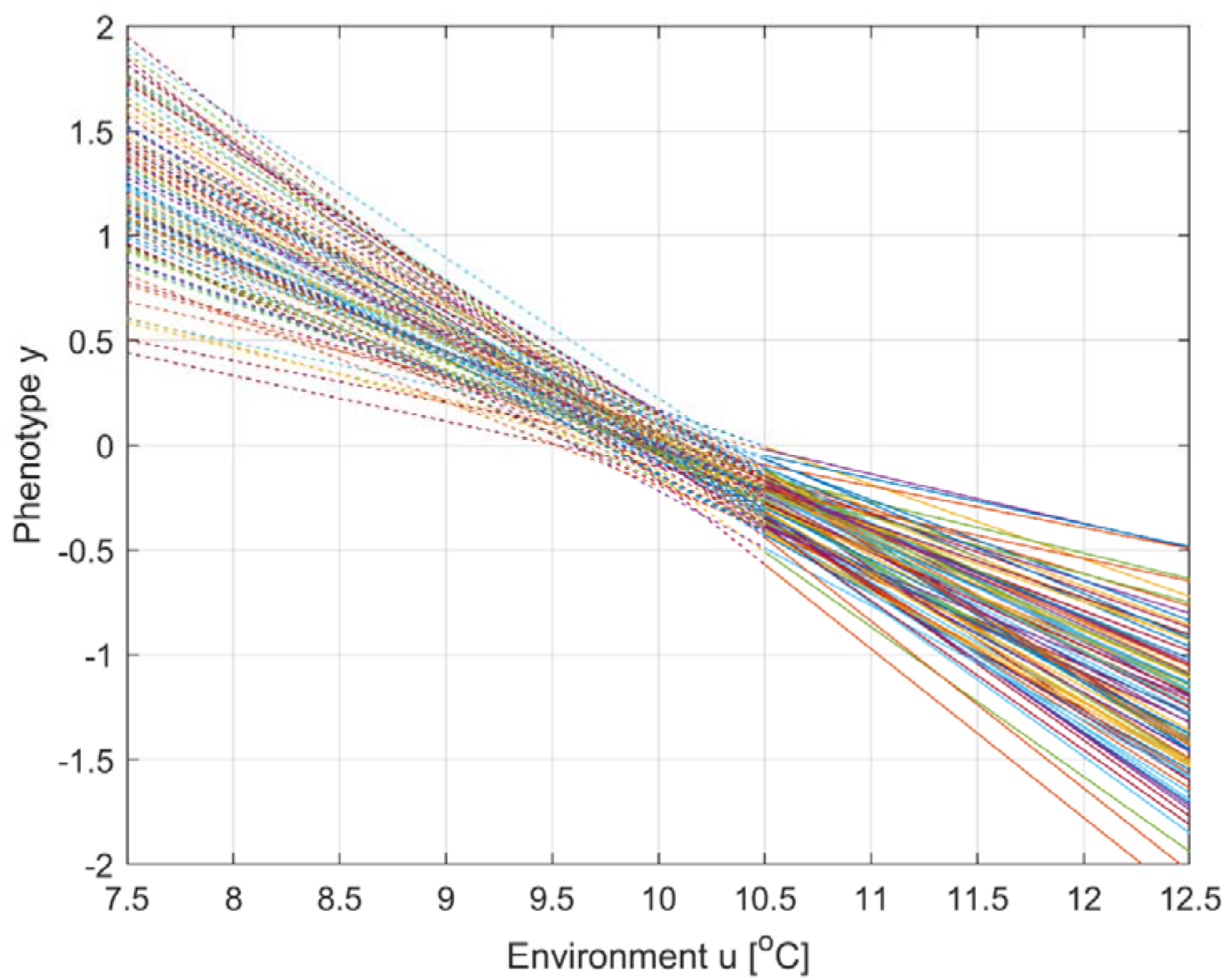
Reaction norms for 100 individuals in a population according to Equation (1), with 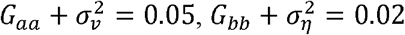 and *G_ab_* = 0. The environmental zero-point is *u_ref_* = 10 °C, which since *G_ab_* = 0 also is the temperature *u*_0_ to which the population is fully adapted. The mean trait values are 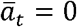 and 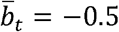. Solid lines indicate the range of data used for parameter estimation and mean trait predictions in simulations. Note that *u_ref_* = *u*_0_ = 10 °C is not within that range.

### 2.3 Mean trait prediction equations

A fundamental equation for mean trait predictions follows from Equation (2a) as

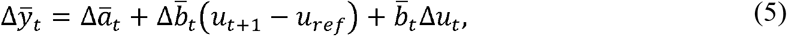

where Δ*u_t_* = *u*_*t*+1_ - *u_t_*, 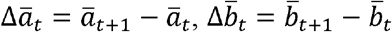 and 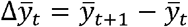 are changes per generation. From this follows that the value of *u_ref_* has nothing to say in special cases with constant phenotypic plasticity slopes, i.e., when 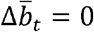. In such cases we simply have 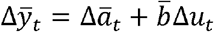, where 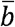 is constant, or only 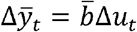 if 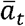 does not evolve.

As shown in Appendix 1, Equation (5) leads to equations for 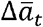 and 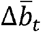 as functions of the phenotypic selection gradient *β_y,t_*,

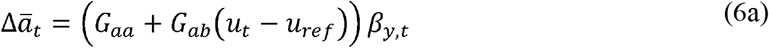

and

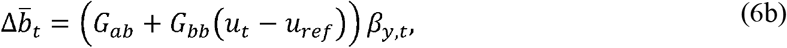

where

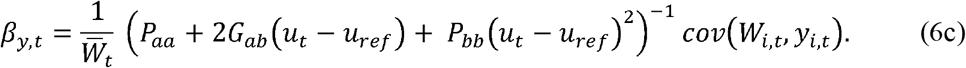

In addition to time series of *u_t_* and *y_i,t_*, we thus need parameter values for *u_ref_, G_aa_, G_ab_, G_bb_*, 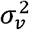 and 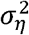, and a time series of individual fitness values *W_i,t_*. For mean trait predictions we also need initial values. Note that these equations are valid only when the genetic relationship matrix is a unity matrix (Ch. 26, Lynch & Walsh, 1998).

### 2.4 Prediction error minimization method

From predicted mean intercept and plasticity slope values found by Equations (6a,b) follow predicted values of 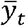 from Equation (2a). The prediction equations can thus be used for parameter estimation in a prediction error minimization method (PEM), as shown in Fig. 2. As follows from Equations (6a,b,c), we can then set *G_aa_* to any value, and estimate *G_ab_, G_bb_*, 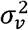 and 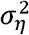 relative to that value.

**Figure 2.**
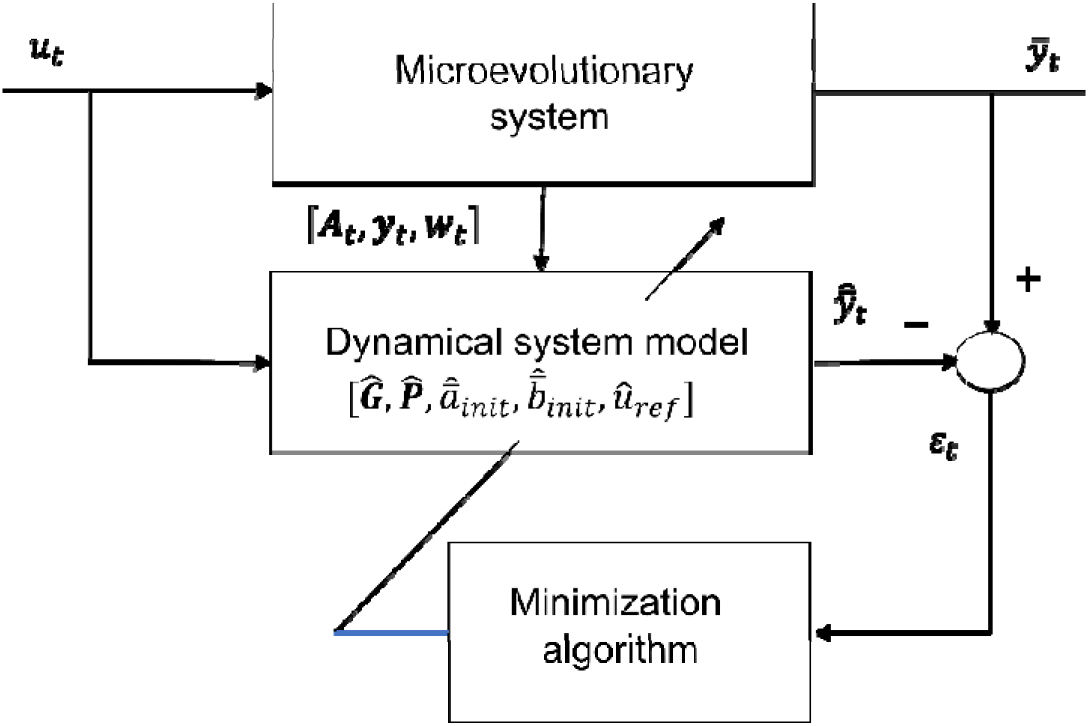
Block diagram of microevolutionary PEM, with dynamical tuning model based on an intercept-slope reaction norm model with mean traits and. Here, and are the known environmental input and the known mean phenotypic value at time, respectively. is the additive genetic relationship matrix, which here is assumed to be, while and are vectors of individual phenotypic and relative fitness values, respectively. The and matrices include the system parameters, while, and are the initial mean trait values and the environmental zero-point, respectively. Assuming data over generations, all these model parameters are tuned until is minimized, with and.

### 2.5 Effects of environmental zero-point errors

With an environmental zero-point 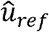 instead of *u_ref_*, predictions based on Equation (2a) can be written

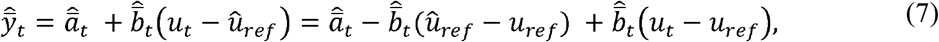

where 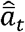 and 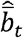 are found from Equations (6a,b) with use of estimated parameter values.

For small values of *G_bb_*, i.e., when *G_bb_* → 0 and *G_ab_* → 0, it follows from Equations (6a,b,c) that 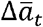 is independent of *u_ref_*, and that 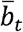 is constant. This results in 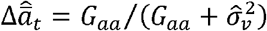, such that only 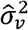 must be tuned in order to minimize 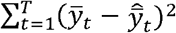. In this case an error in 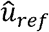 has very little effect on the change in 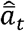 over many generations, as also follows from Equation (5).

For larger values of *G_bb_* and *G_ab_*, 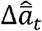 will be affected by an error in 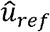, and good predictions 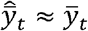 for *t* = 1 to *T* can then only be obtained by parameter tuning such that 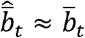 over all generations. That is possible because *u_ref_* appears in both nominator and denominator of eqn (6b). According to Equation (7) we then find 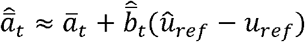, which as shown in Section 3 may result in large errors in predicted changes of 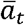 over time.

### 2.6 Effects of modeling errors

Modeling errors will obviously affect predictions of the mean traits. As an example, simulations with the true individual model

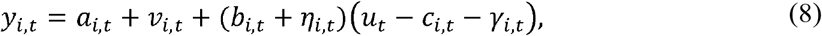

are included in Appendix 2. Here, *c_i,t_* is a perception trait, as discussed in Ergon & Ergon (2017).

## 3 Simulation results

### 3.1 Description of toy example

In the toy example, the environmental input (*u_t_*) is a noisy positive trend in spring temperature, resulting in a noisy negative trend in mean clutch-initiation date 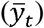 for a certain bird species, approximately as in Fig. 2 in Bowers et al. (2016). The individual phenotypic values are discrete, with days as unit. The individual fitness values (*W_i,t_*) are integers from 0 to 10, with number of fledglings as unit. Generations are assumed to be non-overlapping, and the population size is assumed to be constant. Data for *u_t_, y_i,t_* and *W_i,t_* are generated over 60 generations, where the positive temperature trend begins at generation 10. The population is assumed to be fully adapted to the mean spring temperature 10 °C before generation 10, which is thus the reference environment *u_ref_*, but only data from generations 31 to 60 are used for parameter estimation and mean trait predictions. Note that 10 °C may not be within the range of input data used for parameter estimation (depending on realization). The essential questions are how microevolutionary changes in mean intercept and plasticity slope over generations 31 to 60 can be predicted, and how errors in the estimated or assumed value of *u_ref_* will affect these predictions.

#### 3.1 True model, fitness function, and environmental input signals

Assume that what we consider to be true mean responses, 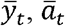 and 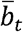, are generated by the state-space model (2a,b). Here, *G_ab_* = 0 in the true system but left as a free parameter in the tuning model in Fig. 2. The individual effects *a_i,t_, b_i,t_, v_i,t_* and *η_i,t_* are at each generation drawn from populations with normal distributions around 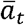, 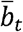, 0 and 0, respectively.

The individual fitness function is assumed to be rounded values of

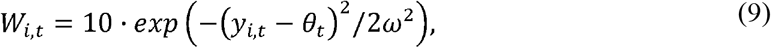

where *θ_t_* is the phenotypic value that maximizes fitness, while *ω*^2^ = 10. The discrete values of *W_i,t_* (number of fledglings) are thus integers from 0 to 10.

Also assume a stationary or slowly varying mean *μ_⋃,t_* of a stochastic environment, with added iid zero mean normal random variations *u_n,t_* with variance 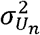, i.e., *u_t_* = *μ_⋃,t_* + *u_n,t_*, and that the population is fully adapted to a stationary stochastic environment with *μ_⋃,t_* = *u_ref_* = *u*_0_ = 10 °C (as in Fig. 1). In a corresponding way assume that *θ_t_* = *μ*_Θ,*t*_ + *θ_n,t_*, where *θ_n,t_* is iid zero mean normal with variance 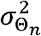, and where *u_n,t_* and *θ_n,t_* are correlated with covariance *σ*_Θ_*n*_⋃_*n*__. Following Lande (2009), we may assume that juvenile birds of generation *t* are exposed to the environment *u_t-τ_* during a critical period of development a fraction of a generation before the adult phenotype is expressed and subjected to natural selection. We will define *θ_t_* = –2(*u_t_* – 10), which implies a linear relationship *μ*_Θ,*t*_ = −2(*μ_⋃,t_* - 10), variances 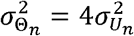, and covariance 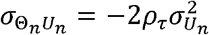, where *ρ_τ_* is the autocorrelation of background environmental fluctuations. We will assume 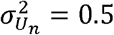 and *ρ_τ_* = 0.25. The optimal value of the mean plasticity slope in a stationary stochastic environment is then 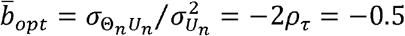 (as in Fig. 1) (Ergon & Ergon, 2017).

Further assume that *u_t_* and *θ_t_* are noisy ramp functions as shown in Fig. 3, with the ramp in *μ_⋃,t_* starting from 10 °C at *t* = 10 generations. The choice of a negative trend in *θ_t_*, and thus in 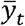, results in earlier clutch-initiation dates as a result of the positive temperature trend.

**Figure 3.**
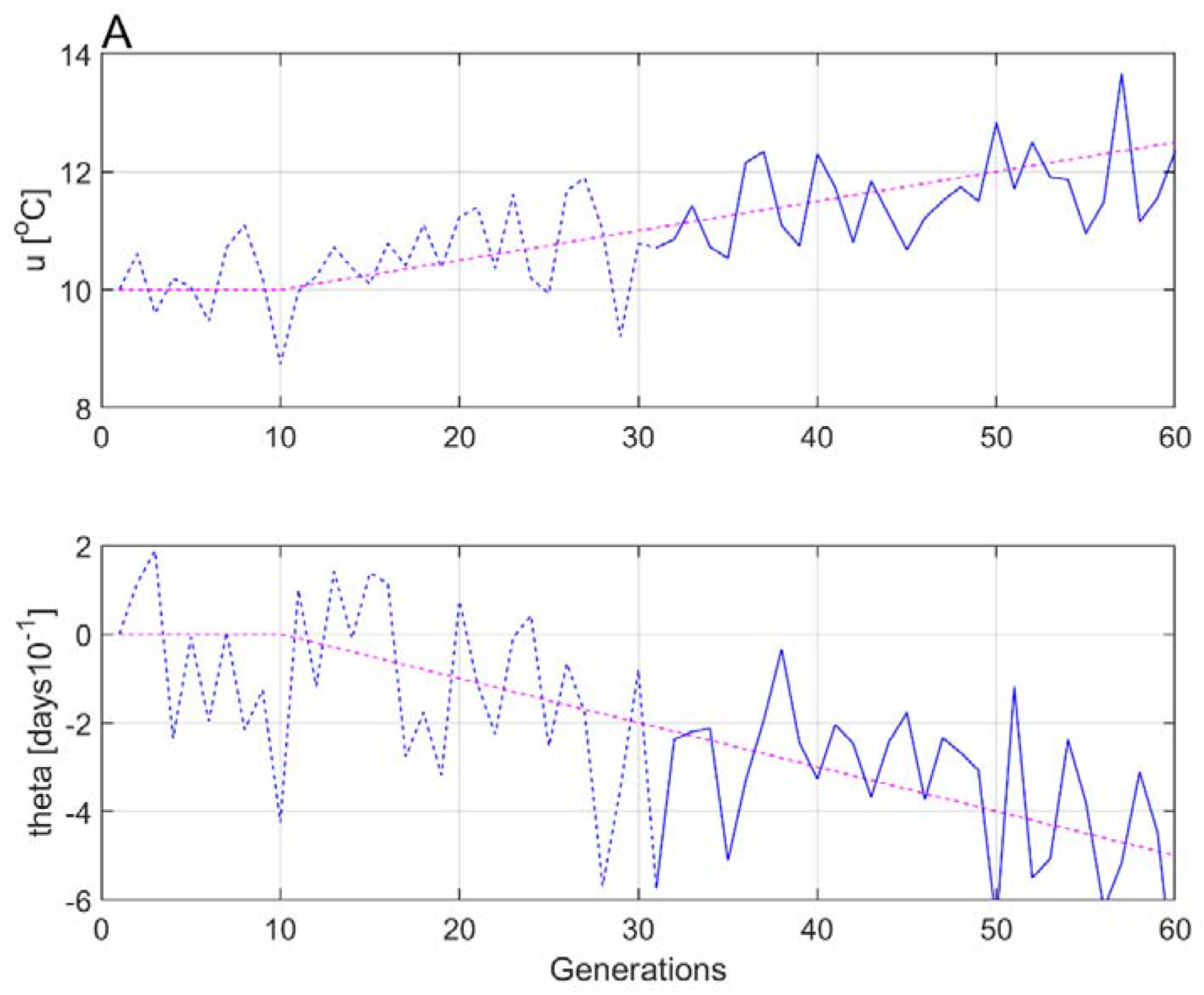
Noisy ramp functions *u_t_* (panel A) and *θ_t_* (panel B), with *u_ref_* = 10, 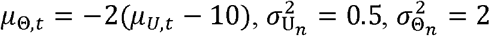 and *σ*_⋃_*n*_,Θ_*n*__ = −0.25. The solid parts of the curves indicate data used for parameter estimation and mean trait predictions (compare with Fig. 1, where reaction norms for *u_t_* > 10.5 are indicated by solid lines).

Fig. 4 shows typical individual phenotypic (clutch-initiation date) and fitness (number of fledglings) values for the true model with population size *N* = 100 at generation 45 in Fig. 3. The figure shows that the most negative (earliest) dates give the highest number of offspring, and the population is thus under directional selection towards earlier clutch-initiation dates. The zero date is the mean clutch-initiation date before the positive temperature trend sets in at generation 10 in Fig. 3, when the population is assumed to be under stabilizing selection and fully adapted to the stationary stochastic temperature.

**Figure 4.**
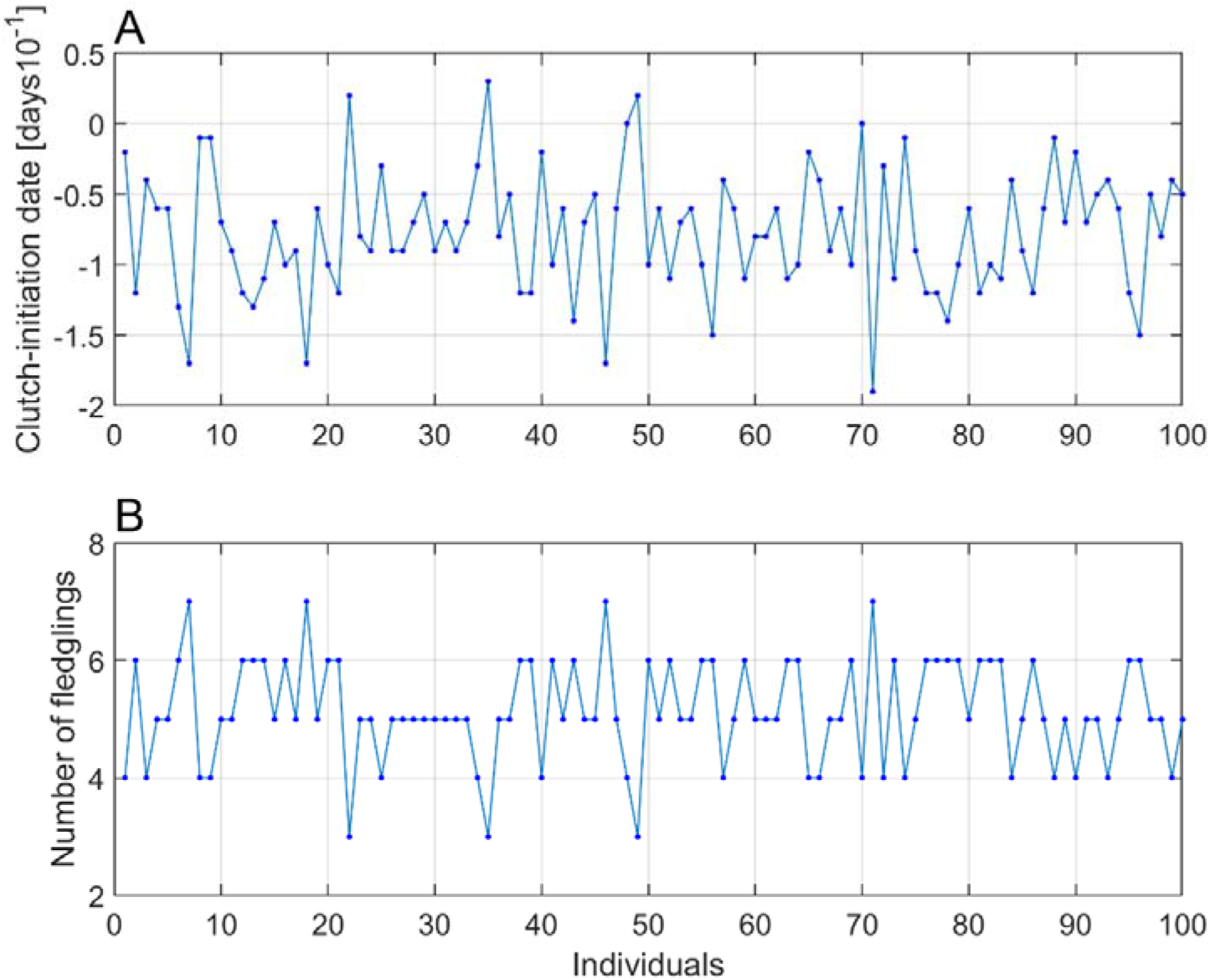
Individual clutch-initiation dates *y_i,t_*, with a range from —19 to 3 days (phenotypic values, Panel A), and number of fledglings *W_i,t_* (fitness, Panel B), at a generation where the population is under directional selection towards earlier clutch-initiation dates. As indicated by dots the number of offspring are integers, while the phenotypic values are integers divided by ten (i.e., days).

#### 3.2 Parameter estimation and mean trait prediction results

Parameter estimation and mean trait prediction results were found by use of the MATLAB function *fmincon* in the PEM method in Fig. 2. Results with use of input-output data from *t* = 31 to 60 with population size *N* = 100 are given in Table 1. The relative errors in total change of predictions over 30 generations are included, computed as 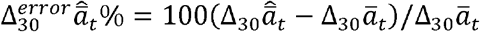 etc., where 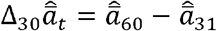 and 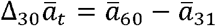. The final values 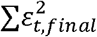 of 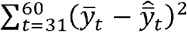 are also included, as they indicate the degree of optimization success. Results are presented as mean values and standard errors, *Mean* ± *SE*, based on 100 repeated simulations with different realizations of random inputs.

Given the model in Equations (2a,b) and (9), there are six parameter values to be estimated (while 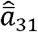 follows from Equation (2a) with 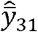 sat to zero). In the optimizations the initial values of 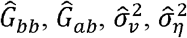 and 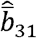 were set to zero, while the initial value of 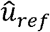 was set to 10 (when 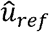 was a free variable). The true value 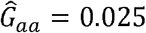 was used, such that estimates of *G_bb_, G_ab_*, 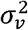 and 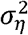 are found relative to *G_aa_* = 0.025. Table 1 presents results for three cases, first for 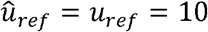 (Case 1), second for 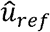 as free variable (Case 2), and third for 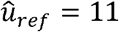 (Case 3), which is the expected initial value in the time series used. The estimates of 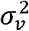 and 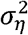 have in all cases rather large standard errors, and in some cases also large bias errors. What is more interesting is that the prediction errors 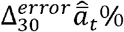 and 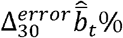 are small in Case 1 (with the correct value 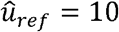). In Case 2 (with 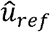 as a free variable), 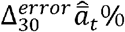 has a large standard error. With 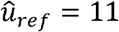 (Case 3), 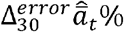 has a large bias error. In this case also the estimates of *G_bb_, G_ab_* and 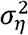 are clearly biased. In all cases 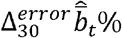 is close to the result with the correct value 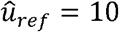 (Case 1), as explained in Subsection 2.5.

Table 1 includes theoretical prediction error results based on Equation (7), 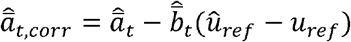. These are in all cases close to the results with 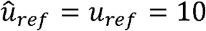 (Case 1).

The results for Case 1 and Case 2 are very much improved with population size *N* = 10,000, while the standard errors in Case 3 were only marginally improved by an increased population size (Appendix 3). The results for 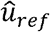 in Case 2 were for example improved from 9.92 ± 0.66 to 9.95 ± 0.19 (ideally 10).

**Table 1.**
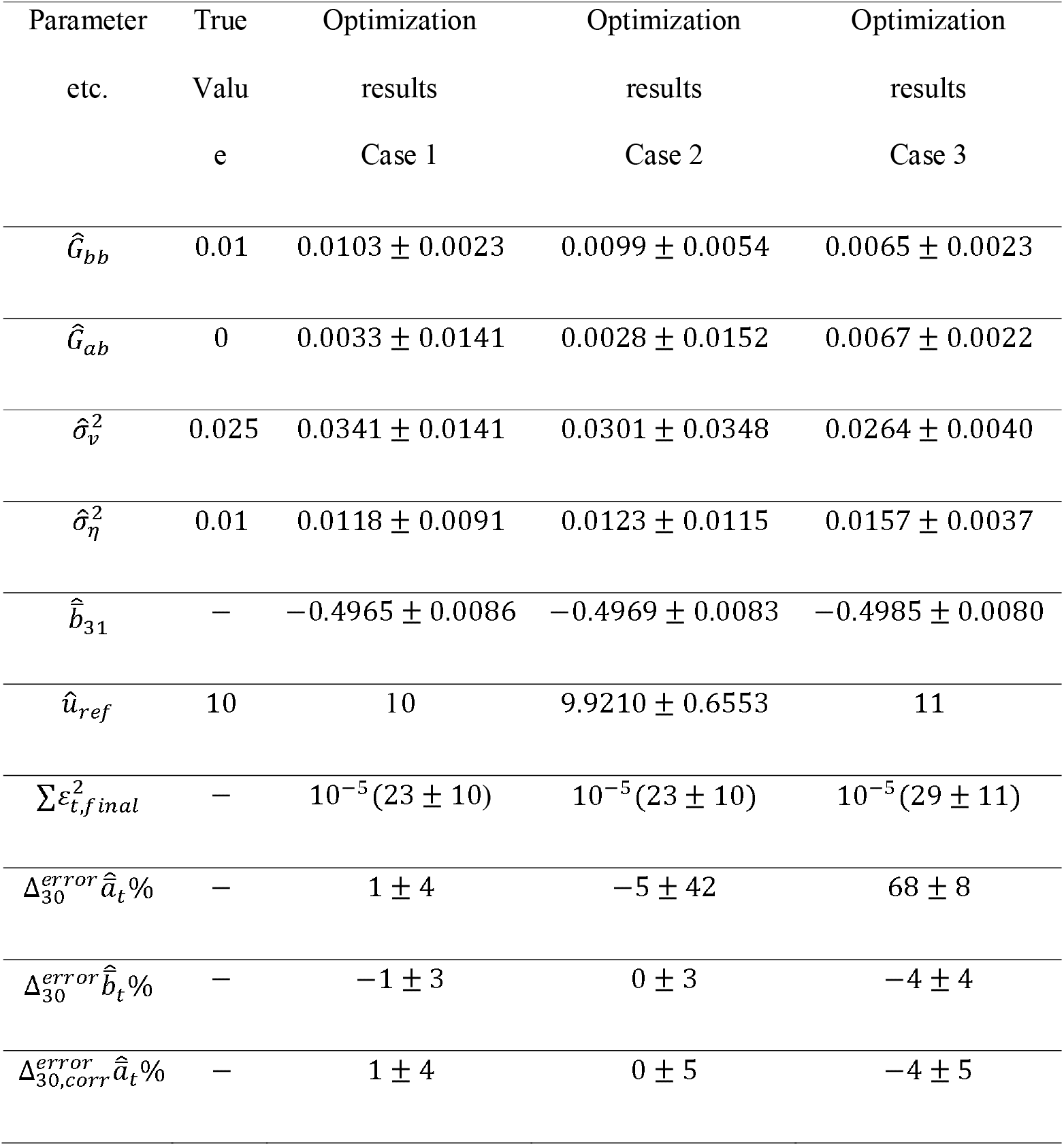
Estimation and prediction results with true system responses generated by means of Equations (2a,b) and (9). Results are for cases with population size *N* = 100 and perfect observations of *y_i,t_* and *W_i,t_,* and they are based on 100 simulations with different realizations of all random input variables. Case 1: 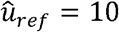 (the true value). Case 2: 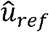 as a free variable. Case 3: 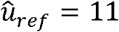 (expected initial value in optimization data). Here, 9% of the simulations were discarded because 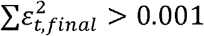.

As shown in Table 1, a large error in the assumed environmental zero-point 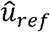 results in large errors in predicted changes of in 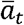 over 30 generations (Case 3). Table 2 shows prediction results for more moderate errors in 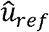, as well as for 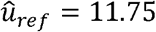 (expected mean value in optimization data), and they are all in accordance with Equation (7).

**Table 2.** Errors in predicted total relative change of 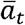 and 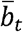 over 30 generations, as functions of the environmental zero-point 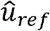 used in the optimization procedure.

| $\hat{u}_{ref}$ | $\Delta_{30}^{error} \hat{a}_t \%$ | $\Delta_{30}^{error} \hat{b}_t \%$ |
| --- | --- | --- |
| 9.75 | $-16 \pm 4$ | $0 \pm 3$ |
| 10 | $1 \pm 4$ | $-1 \pm 3$ |
| 10.25 | $17 \pm 4$ | $0 \pm 3$ |
| 10.5 | $34 \pm 6$ | $-2 \pm 3$ |
| 11 | $68 \pm 8$ | $-4 \pm 4$ |
| 11.75 | $115 \pm 14$ | $-8 \pm 6$ |

Fig. 5 shows predicted mean values 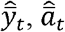 and 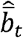, as compared to true mean values 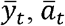 and 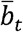, for Case 1 and Case 3 in Table 1.

**Figure 5.**
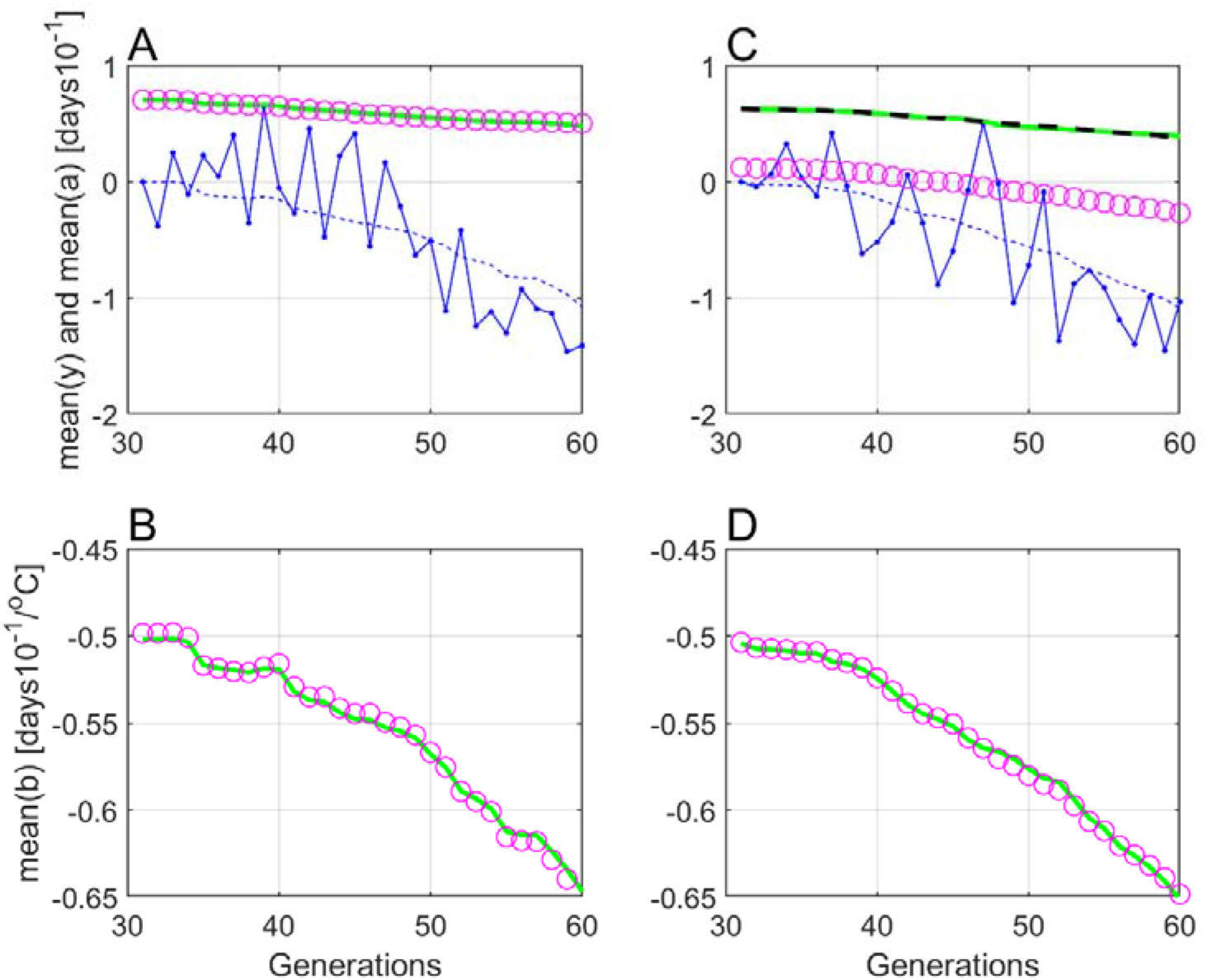
Typical responses for Case 1 and Case 3 in Table 1. True 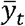 values are shown by solid blue lines. All parameter values except 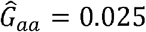 were initially set to zero, which gave predictions 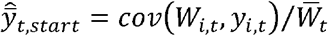 as shown by dashed blue lines. Final predictions 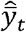 are shown by blue dots. True 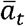 and 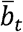 responses are shown by green lines, while predictions 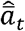 and 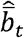 are shown by magenta circles. Panels A and B show results for Case 1 with 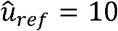 (true value), while panels C and D show results for Case 3 with 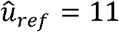. Here, the theoretical predictions 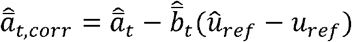 are included as black dashed line. Note that 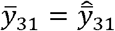 is set to zero, such that 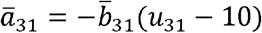 and 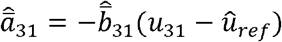, where *u*_31_ is not quite the same from realization to realization.

### 4 Discussion

It is well documented that populations adjust to climate change by means of individual plasticity, but few reports on adaptation by means of genetically based microevolution caused by phenotypic selection (Merilä & Hendry, 2014). The main point in this article is that disentanglement of these separate effects requires that the environmental zero-point *u_ref_* is defined, and that this should not be done arbitrarily. Instead, it should be based on the environment *u*_0_ where the phenotypic variance is at a minimum (Definition 1 and Fig. 1). This definition can be extended to multivariate cases. As shown in a toy example, large errors in *u_ref_* may lead to large errors in predicted microevolutionary changes over time (Table 1 and Fig. 5). Such large errors in *u_ref_* may occur when the range of environmental data used for predictions is far from the mean value of the stochastic environment the population is adapted to.

Although the plastic response to climate change is a result of individual plasticity, it should be noted that individual traits do not determine the environmental value *u*_0_ in Definition 1, but instead the phenotypic variance (Fig. 1). Similarly, individual traits not enter into the prediction equations (6a,b), but instead the individual phenotypic and fitness values (Fig. 4).

In theoretical studies it is often assumed that the environmental variable is scaled such that *u_ref_* = *u*_0_ = 0 (Lande, 2009; Chevin & Lande, 2015). This can be done also in databased applications, provided that *u*_0_ is known, and that the correction term in Definition 1 is zero.

The toy example used in the simulations is a simplification of reality, because changing spring temperatures affect fitness in a complex way (Bowers et al., 2016). It still suffices to show that the environmental zero-point together with initial mean trait and parameter values can be estimated from environmental, phenotypic and fitness data, by use of the prediction error minimization method in Fig. 2. The simulations make use of an environmental trend, as in noisy temperature trends caused by climate change (Fig. 3), and a correct environmental zero-point then results in quite good predictions of changes in mean quantitative traits over time (Table 1, Case 1). Although these predictions are based on parameter estimation, all the separate parameter estimates as such are not especially good, although they were considerably improved when the population size was increased from 100 to 10,000 (Appendix 3). The environmental zero-point can also be estimated, but with large standard errors, especially for small population sizes, and this results in a correspondingly large standard error in predicted change in mean intercept 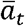 over time (Table 1, Case 2). An estimated zero-point may still be a better choice than use of an initial environmental value in a recorded time series, or the mean value, which may give large errors in predicted changes in mean quantitative traits (Table 1, Case 3). Another alternative may be to use the mean value of a past stationary stochastic environment, which the population is judged to have been fully adapted to.

It is here assumed that the genetic relationship matrix is an identity matrix, and the simulation results are obtained by use of a prediction error minimization method. However, the fact that errors in the environmental zero-point may cause large errors in predictions of microevolution, as discussed in Subsection 2.5, is a generic problem. Independent of prediction method and the complexity of the model, an error in the environmental zero-point implies that an erroneous model is fitted to the input-output data, and that must inevitably result in prediction errors. An alternative view when 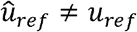 is constant (Case 3), is that the tuning model in Fig. 2 still uses the correct value of *u_ref_*, but then with an error term 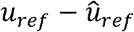 added to the environmental input. In order to minimize 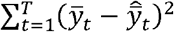, this input error must as good as possible be compensated by errors in estimated parameter values, resulting in prediction errors. Note that this argument is independent of the specific parametrizations used in the microevolutionary and tuning models in Fig. 2. It is in any case no reason to believe that prediction errors caused by 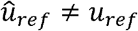 will disappear in cases where the genetic relationship matrix is not a unity matrix, and when other parameter estimation and mean trait prediction methods are used. A more specific argument regarding BLUP/REML parameter estimation is given in Appendix 4. It must thus be expected that predictions of microevolutionary changes over time depend on the chosen environmental zero-point (reference environment), and such predictions cannot therefore be trusted unless the chosen environmental zero-point can be trusted. Exceptions are here cases with a constant mean plasticity slope, where the change in mean reaction norm intercept per generation according to Equation (5) is independent of the environmental zero-point. This implies that a nearly constant mean plasticity slope must be expected to result in small errors in the predicted changes of the mean intercept, also if there is an error in the environmental zero-point.

## Supporting information

MATLAB code

## Acknowledgement

The derivation of prediction equations in Appendix 1 was inspired by a review comment by Michael Morrissey, although on a quite different manuscript with a derivation directly based on the Price equation. I thank University of South-Eastern Norway for support and funding.

## Author Contribution

Rolf Ergon is the sole author of this article.

## Data Accessibility Statement

MATLAB code for simulations is given in Supplementary Material archived in *bioRxiv*, **doi:** https://doi.org/10.1101/2022.01.07.475361

## Conflict of interest

The author declares no conflict of interest.

## Appendix 1. Derivation of prediction equations

The goal is to find Equations (6a,b) for incremental changes 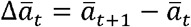 and 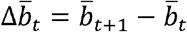, and thus also 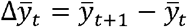, by means of a common selection gradient *β_y,t_*. Note that these equations implicitly assume that the additive genetic relationship matrix ***A_t_*** in Fig. 2 is a unity matrix, if not the genetic relationships will simply be ignored.

As stated in the main text, a fundamental equation for mean trait predictions follows from Equation (2a) as

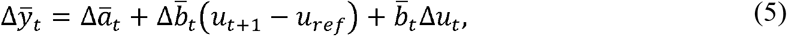

where Δ*u_t_* = *u*_*t*+1_ – *u_t_*. From this follows that the value of *u_ref_* has nothing to say in special cases with constant mean plasticity slopes, i.e., when 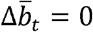. In such cases we simply have 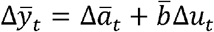, where 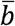 is constant, or only 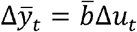 if 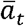 does not evolve.

In the general case, 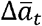 and 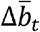 are found from Equations (6a,b), as derived below. Assuming a constant environment, i.e., for Δ*u_t_* = 0, Equation (5) gives 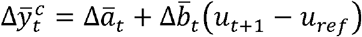, which by means of the breeder’s equation (Lande, 1979) also can be written as

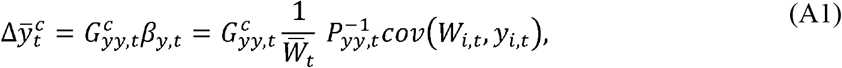

where *β_y,t_* is the selection gradient, while 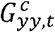 and *P_yy,t_* are additive genetic and phenotypic variances, respectively. Here, *W_i,t_* is the individual fitness, with population mean value 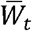, while *y_i,t_* is the individual phenotypic value. We must assume that values of *W_i,t_* and *y_i,t_* are available.

In order to find expressions for 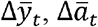 and 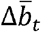, we may express 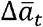 and 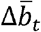 by means of *β_y,t_* according to

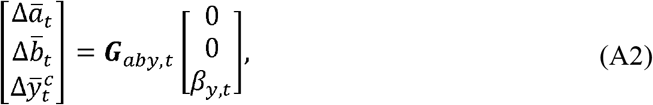

where the *G_aby,t_* matrix is found via the linear transformation of the vector [*a_i,t_ b_i,t_*]^*T*^ onto the vector [*a_i,t_ b_i,t_ y_i,t_*]^*T*^. By use of Equation (1) we find (using 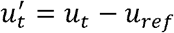)

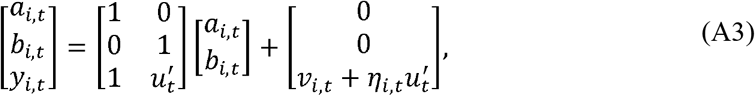

such that the additive genetic covariance matrix of [*a_i,t_ b_i,t_ y_i,t_*]^*T*^ specific to the environment 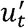 becomes

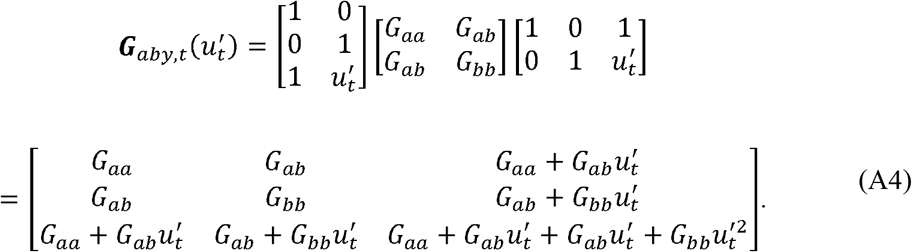

From Equations (A1), (A3) and (A4) follow

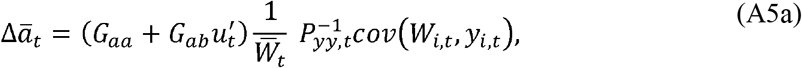

and

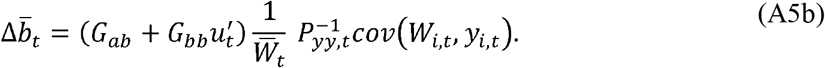

where

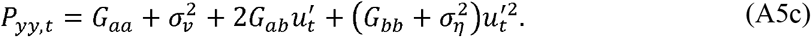

Since only a single environmental value is involved in Equations (A5a,b), they are valid in any environment, also if it varies with time, and 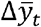 thus follows by inserting 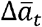 and 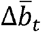 from Equations (5a,b) into Equation (5). For computations of 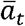 and 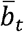, and thus 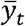, initial values of 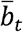 and 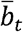 are needed.

Alternatively, 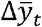 is found as

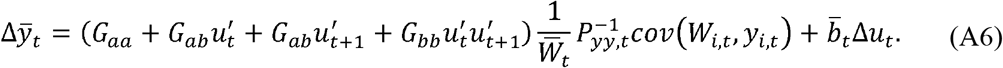

Note that 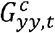 in Equation (A1) follows from *G_yy,t_* according to Equation (A6) by setting 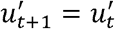, i.e., under the assumption of a constant environment. The same expression for 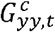 also follows from Equation (A4).

## Appendix 2. Modeling error results

Simulations with a true three-trait reaction norm model according to Equation (8), i.e., with a perception trait in addition to the intercept and plasticity traits (Ergon & Ergon, 2017), but with a two-trait tuning model, gave increased prediction errors as compared to results in Table 1. The population size was *N* = 100. Simulations with *G_cc_* = *G_aa_* = 0.025 and the true value *u_ref_* = 0, gave the following results: With *G_ac_* = 0 the prediction errors were 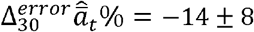 and 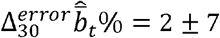. With *G_ac_* = –0.01 the results were 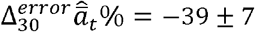 and 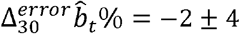, while *G_ac_* = 0.01 gave 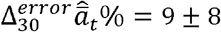 and 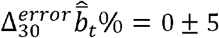. Note that 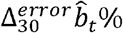 is very much the same for all values of *G_ac_*.

Typical responses are shown in Fig. A1. Note especially that the change in 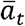 over time for *G_ac_* ≤ 0 is underestimated, which indicates that modeling errors in general may result in underestimated changes in 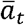. This should be compared with the results for Case 3 in Table 1, where changes in 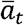 are overestimated.

**Figure A1.**
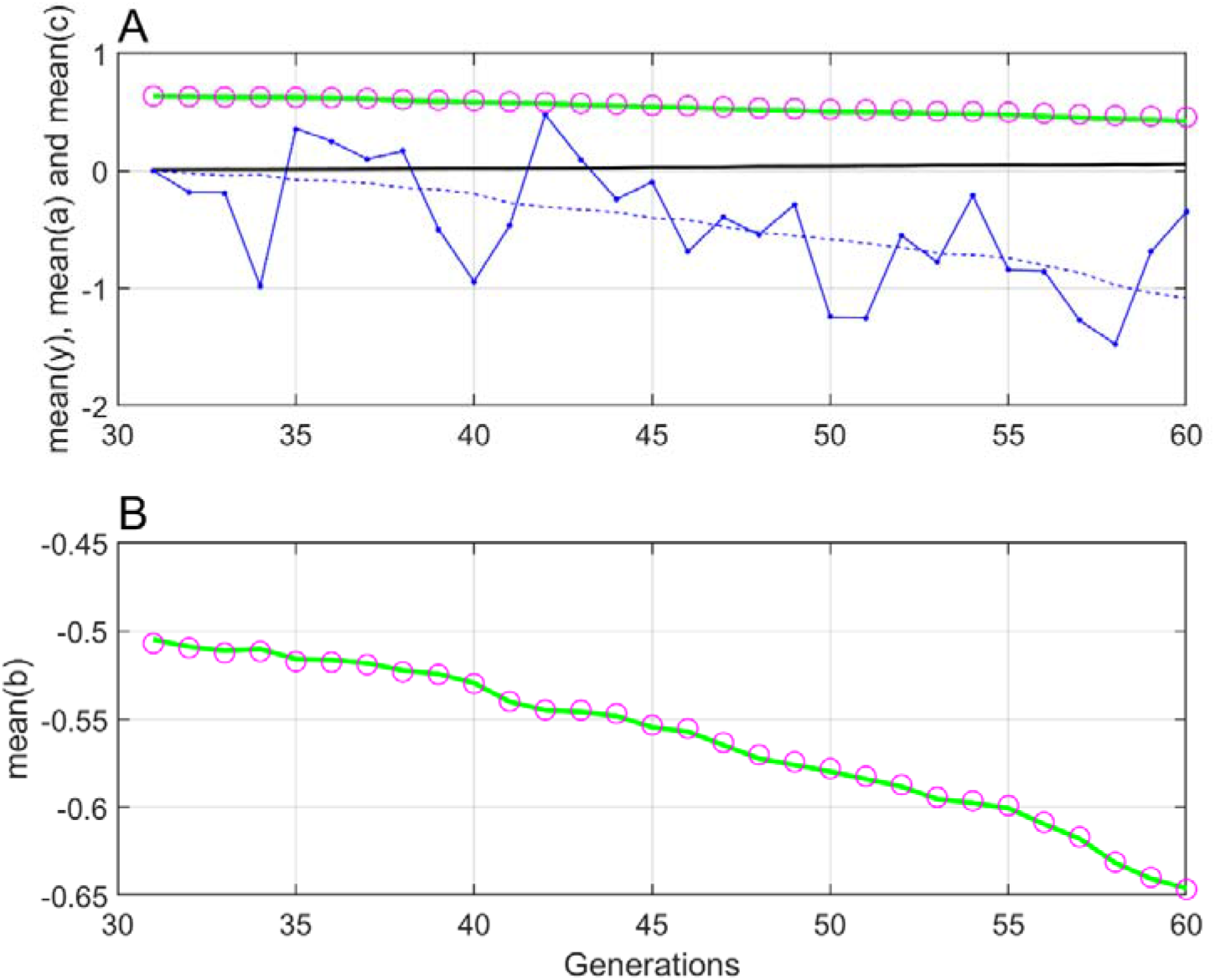
Typical responses as for Case 1 in Fig. 5, but with a true three-trait model according to Equation (8) with *G_cc_* = *G_aa_* = 0.025 and *G_ac_* = 0. The true 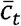 response is shown by black line.

## Appendix 3. Results with increased population size

Table A1 shows results as in Table 1, but with population size *N* = 10,000.

**Table A1.** Estimation and prediction results as in Table 1, but with population size *N* = 10,000. In Case 3, 16% of the simulations were discarded because 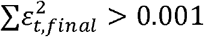.

| Parameter<br>etc. | True<br>value | Optimization<br>results<br>Case 1 | Optimization<br>results<br>Case 2 | Optimization<br>results<br>Case 3 |
| --- | --- | --- | --- | --- |
| $\hat{G}_{bb}$ | 0.01 | $0.0100 \pm 0.0003$ | $0.0103 \pm 0.0023$ | $0.0064 \pm 0.0017$ |
| $\hat{G}_{ab}$ | 0 | $0.0006 \pm 0.0033$ | $0.0002 \pm 0.0029$ | $0.0068 \pm 0.0015$ |
| $\hat{\sigma}_v^2$ | 0.025 | $0.0269 \pm 0.0064$ | $0.0277 \pm 0.0146$ | $0.0256 \pm 0.0034$ |
| $\hat{\sigma}_\eta^2$ | 0.01 | $0.0103 \pm 0.0021$ | $0.0102 \pm 0.0026$ | $0.0161 \pm 0.0027$ |
| $\hat{b}_{31}$ | — | $-0.4958 \pm 0.0068$ | $-0.4962 \pm 0.0086$ | $-0.4969 \pm 0.0089$ |
| $\hat{u}_{ref}$ | 10 | 10 | $9.9483 \pm 0.1921$ | 11 |
| $\sum \varepsilon_{t,final}^2$ | — | $10^{-5}(0 \pm 2)$ | $10^{-5}(1 \pm 2)$ | $10^{-5}(10 \pm 10)$ |
| $\Delta_{30}^{error} \hat{a}_t \%$ | — | $0 \pm 1$ | $-3 \pm 12$ | $68 \pm 6$ |
| $\Delta_{30}^{error} \hat{b}_t \%$ | — | $0 \pm 1$ | $0 \pm 1$ | $-4 \pm 3$ |
| $\Delta_{30,corr}^{error} \hat{a}_t \%$ | — | $0 \pm 1$ | $0 \pm 1$ | $4 \pm 4$ |

## Appendix 4. REML parameter estimation

The reaction norm model in Equation (1) can by use of Equation (2a) be written as

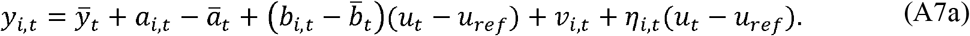

From this follows the random mixed model for the population,

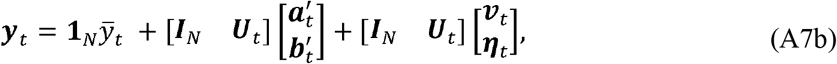

where 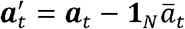 and 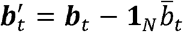 (where **1**_N_ is an *N* × 1 vector of ones). Here, ***y**_t_, **a**_t_, 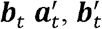, **v**_t_* and ***η**_t_* are *N* × 1 vectors of individual values at time *t*, while 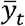 is the scalar valued fixed effect. The environmental input matrix is ***U**_t_* = (*u_t_* – *u_ref_*)***I**_N_*, where ***I**_N_* is the *N* × *N* unity matrix. The expected values 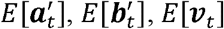 and *E*[***η**_t_*] are all zero by definition (Ch. 26, Lynch and Walsh, 1998).

The starting point for REML parameter estimation based on the model (A7b) is the phenotypic population covariance matrix 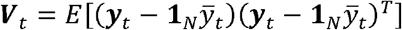 (Ch. 27, Lynch and Walsh, 1998). For the model in Equation (1) this covariance matrix is

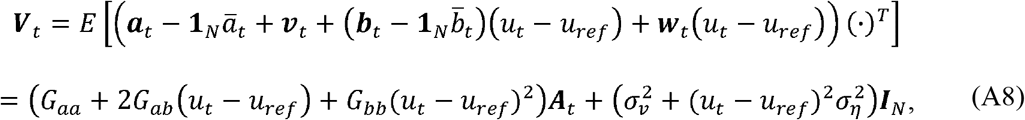

where ***A**_t_* is the genetic relationship matrix (Ch. 27, Lynch and Walsh, 1998). From Equation (A8) follows that REML parameter estimation results depend on *u_ref_*, such that errors in the environmental zero-point give errors in estimated parameter values. As in parameter estimation based on Equations (6a,b), we can in REML estimation set *G_aa_* to any value, and estimate other parameters in relation to that. It follows from Equation (A8) that we cannot find 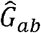 separate from 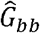, and 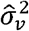 separate from 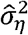, unless data for several different values of *u_t_* are used. This may be compared with the PEM method in Fig. 2, as used in the simulations, where data from all available generations are utilized.

## Notes

### Competing Interest Statement

The authors have declared no competing interest.

### Summary of Updates

1. Fig. 2 caption will hopefully be converted correctly. 2. MATLAB code is now complete.

## References

Chevin, L.-M., & Lande, R. (2015). Evolution of environmental cues for phenotypic plasticity. Evolution, 46(2), 390–411. doi:10.1111/evo.12755

Bowers, E.K., Grindstaff, J.L., Soukup, S.S., Drilling, N.E., Eckerle, K.P., Sakaluk, S.K., & Thompson, C.F. (2016). Spring temperatures influence selection on breeding date and the potential for phenological mismatch in a migratory bird. Ecology, 97(10), 2880–2891.

Ergon, T., & Ergon, R. (2017). When three traits make a line: evolution of phenotypic plasticity and genetic assimilation through linear reaction norms in stochastic environments. J. Evol. Biol. 30, 486–500. doi: 10.1111/jeb.13003

Ergon, R. (2018). The environmental zero-point problem in evolutionary reaction norm modeling. Ecol Evol. 8, 4031–4041. doi: 10.1002/ece3.3929

Fossen, E.I.F., Pélabon, C. & Einum, S. (2018). An empirical test for a zone of canalization in thermal reaction norms J. Evol. Biol. 31, 936–943. doi: 10.1111/jeb.13287.

Lande, R. (1979). Quantitative genetic analysis of multivariate evolution, applied to brain:body size allometry. Evolution 33, 402–416.

Lande, R. (2009). Adaptation to an extraordinary environment by evolution of phenotypic plasticity and genetic assimilation. J. Evol. Biol. 22, 1435–1446. doi: 10.1111/j.1420-9101.2009.01754.x

Ljung, L. (2002). Prediction Error Estimation Methods. Circuits, Systems, and Signal Processing 21/1, 11–21.

Lynch, M., & Walsh, B. (1998). Genetics and Analysis of Quantitatve Traits. Sinauer Associates, Mass.

Merilä, J., & Hendry, A.P. (2013). Climate change, adaptation, and phenotypic plasticity: the problem and the evidence. Evol. Appl. 7, 1–14. doi: 10.1111/eva.12137.

Morrissey, M.B., Kruuk, L.E.B, & Wilson, A.J. (2010) The danger of applying the breeder’s equation in observational studies of natural populations. J. Evol. Biol. 23, 2277–2288. doi: 0.1111/j.1420-9101.2010.02084.x

