## Supplementary material for "The important choice of reference environment in microevolutionary climate response predictions": MATLAB code

### MATLAB code for

### The important choice of reference environment

### in microevolutionary climate response predictions

```
%----------------------------------------------------------
% Parameter estimation of intercept-slope model with
% fmincon() in Matlab.

% Rolf Ergon
% University of South-Eastern Norway
% February 23, 2022
%----------------------------------------------------------

clear

M=1
countW=0;       	% count realizations with mean(W0)=0
countfval=0;    	% count realizations with good convergence

for m=1:M
    m
T=60;
N=100;
wsquare=10;
vartheta=2;
varu=0.5;
rho=0.25;
Gaa=0.025;
vare=Gaa;
Gab=0;
Gbb=0.01;
vareta=Gbb ;
Paa=Gaa+vare;
Pab=Gab;
Pbb=Gbb+vareta;
G=[Gaa Gab ; Gab Gbb];
P=[Paa Pab ; Pab Pbb];

%% Generate u and theta sequences
uplot=10*ones(1,T);
u2=zeros(1,T);
du=zeros(1,T);
dtheta=zeros(1,T);
for i=2:T
    du(i)=sqrt(varu)*randn;
    dtheta(i)=du(i)*rho*sqrt(vartheta/(varu))+sqrt(vartheta*(1-rho^2))*randn;
    if i>10
        uplot(i)=10+(i-10)/20;
    end
end
u=uplot+du;
theta=-2*(uplot-10)+dtheta;

%% Individual population traits around abar, bbar1 and bbar2
for t=1:T
    a(:,t)=sqrt(Gaa)*randn(N,1);
    b(:,t)=Gab*a(:,t)/Gaa+sqrt(Gbb-Gab^2/Gaa)*randn(N,1);
    v(:,t)=sqrt(vare)*randn(N,1);
    eta(:,t)=sqrt(vareta)*randn(N,1);
    a(:,t)=a(:,t)-mean(a(:,t));
    b(:,t)=b(:,t)-mean(b(:,t));
    v(:,t)=v(:,t)-mean(v(:,t));
    eta(:,t)=eta(:,t)-mean(eta(:,t));
end

%% Simulation of true system
% bbar=-0.25*ones(1,T);
bbar0=-0.5*ones(1,T);
ybar0=0*ones(1,T);
for t=1:T-1
    abar0(31)=ybar0(31)-bbar0(31)*(u(31)-10);
    bterm=(bbar0(t)+b(:,t)+eta(:,t)).*(u(t)-10);
    Y0(:,t)=abar0(t)+a(:,t)+v(:,t)+bterm;
    Y0(:,t)=round(10*Y0(:,t));
    Y0(:,t)=Y0(:,t)/10;
    W0(:,t)=10*exp(-(Y0(:,t)-theta(t)).^2/(2*wsquare));
    W0(:,t)=round(W0(:,t));
    if mean(W0(:,t))==0
        countW=countW+1;		% Count realizations with mean(W0(:,t))==0
        W0(:,t)=0.000001;
    end
    Wbar0(t)=mean(W0(:,t));
    covabW=(N-1)*cov([a(:,t)+v(:,t) b(:,t)+eta(:,t) W0(:,t)])/N;
    covaW(t)=covabW(1,3);
    covbW(t)=covabW(2,3);
    xbar(:,t)=[abar0(t) bbar0(t)]';
    xbar(:,t+1)=xbar(:,t)+G*pinv(P)*[covabW(1,3) covabW(2,3)]'/Wbar0(t);
    xbarny=xbar(:,t+1);
    abar0(t+1)=xbarny(1);
    bbar0(t+1)=xbarny(2);
end
ybar0=mean(Y0);

%% Contraints
Gbb_min=0;Gbb_max=0.1;
Gab_min=-0.1;Gab_max=0.1;
vare_min=0;vare_max=1;
varw_min=0;varw_max=0.1;
bbarinit_min=-1; bbarinit_max=0;
uref_min=9.999999;uref_max=10.000001;
% uref_min=9;uref_max=11;
% uref_min=10.999999;uref_max=11.000001;

par_lb=[Gbb_min,Gab_min,vare_min,varw_min,bbarinit_min,uref_min];
par_ub=[Gbb_max,Gab_max,vare_max,varw_max,bbarinit_max,uref_max];

par_guess=[0 0 0 0 0 10];
par_known=Gaa;
ybarhat_init=0;

%% fmincon

u=u(31:60);
Y=Y0(:,31:59);
abar=abar0(31:60)-ybar0(31);
bbar=bbar0(31:60);
ybar=ybar0(31:59)-ybar0(31);
W=W0(:,31:59);
Wbar=mean(W);

Aineq=[]; Bineq=[]; Aeq=[]; Beq=[];
fun_objective_Example_handle=...
    @(par)fun_objective_Example(par,par_known,u,Y,ybar,W,Wbar,ybarhat_init,N,T);
fun_constraints_Example_handle=...
    @(par)fun_constraints_Example(par,par_known,u,Y,ybar,W,Wbar,ybarhat_init,N,T);
[par_opt,fval,exitflag,output,lambda,grad,hessian] =...
fmincon(fun_objective_Example_handle,par_guess,Aineq,Bineq,Aeq,Beq,par_lb,par_ub,fun_constraints_Example_handle);

%% Simulation with initial model
par_guess=zeros(1,6);
Gbbinit=par_guess(1);
Gabinit=par_guess(2);
vareinit=par_guess(3);
varwinit=par_guess(4);
bbarinit=par_guess(5);
bbarhat=bbarinit*ones(30,1);
ybarhatinit=ybar(1)*ones(30,1);
for t=1:29
    abarhat(1)=ybar(1)-bbarhat(1)*u(1);
    Pyy=Gaa+vareinit+2*Gabinit*u(t)+u(t)^2*(Gbbinit+varwinit);
    covWy=(N-1)*cov(W(:,t),Y(:,t))/N;
    beta(t)=pinv(Pyy)*covWy(1,2)/(1*Wbar(t));
    abarhat(t+1)=abarhat(t)+(Gaa+Gabinit*u(t))*pinv(Pyy)*covWy(1,2)/(1*Wbar(t));
    bbarhat(t+1)=bbarhat(t)+(Gabinit+Gbbinit*u(t))*pinv(Pyy)*covWy(1,2)/(1*Wbar(t));
    ybarhatinit(t+1)=abarhat(t+1)+bbarhat(t+1)*u(t+1);
end

%% Simulation with adapted model
Gbbopt=par_opt(1);
Gabopt=par_opt(2);
vareopt=par_opt(3);
varwopt=par_opt(4);
bbaropt=par_opt(5);
urefopt=par_opt(6);

% fval;

if fval<0.001
    countfval=countfval+1;		% Count useful realizations
    Fvalresult(countfval)=fval;
    Gbboptresult(countfval)=Gbbopt;
    Gaboptresult(countfval)=Gabopt;
    vareoptresult(countfval)=vareopt;
    varwoptresult(countfval)=vareta;
    bbaroptresult(countfval)=bbaropt;
    urefoptresult(countfval)=urefopt;
    opt_resultsresult(countfval,:)=[par_opt Gaa+vareopt+Gbbopt*(urefopt-10)^2];
end

uopt=u-urefopt;

bbarhat=bbaropt*ones(30,1);
ybarhat=0*ones(30,1);
for t=1:29
    abarhat(1)=ybar(1)-bbarhat(1)*uopt(1);
    Pyy=Gaa+vareopt+2*Gabopt*uopt(t)+uopt(t)^2*(Gbbopt+varwopt);
    covWy=(N-1)*cov(W(:,t),Y(:,t))/N;
    betay=inv(Pyy)*covWy(1,2)/Wbar(t);
    abarhat(t+1)=abarhat(t)+(Gaa+Gabopt*uopt(t))*betay;
    bbarhat(t+1)=bbarhat(t)+(Gabopt+Gbbopt*uopt(t))*betay;
    ybarhat(t+1)=abarhat(t+1)+bbarhat(t+1)*uopt(t+1);
end

if fval<0.001
Totalabar=abar(1)-abar(30);
Totalabarhat=abarhat(1)-abarhat(30);
Rel_total_abar_error(countfval)=(Totalabarhat-Totalabar)/Totalabar;
Totalbbar=bbar(1)-bbar(30);
Totalbbarhat=bbarhat(1)-bbarhat(30);
Rel_total_bbar_error(countfval)=(Totalbbarhat-Totalbbar)/Totalbbar;
Totalabarhatcorr=abarhat(1)-bbarhat(1)*(urefopt-10)-abarhat(30)+bbarhat(30)*(urefopt-10);
Rel_total_abar_error_marked(countfval)=(Totalabarhatcorr-Totalabar)/Totalabar;
end

end

%% Plots:
figure(4)
subplot(2,1,1)
plot(Y(:,15)), hold on, grid
plot(Y(:,15),'.b'), hold off
ylabel('Clutch-initiation date [days10^-^1]')
axis([0 100 -2 0.5])
text(0,0.7,'A','FontSize',14)

subplot(2,1,2)
plot(W(:,15)), hold on, grid
plot(W(:,15),'.b'), hold off
axis([0 100 2 8])
text(0,8.5,'B','FontSize',14)
xlabel('Individuals')
ylabel('Number of fledglings')

ybar=[ybar ybarhat(30)];
ybar=[NaN*ones(1,30) ybar];
ybarhat=[NaN*ones(1,30) ybarhat'];
ybarhatinit=[NaN*ones(1,30) ybarhatinit'];
abar=[NaN*ones(1,30) abar];
abarhat=[NaN*ones(1,30) abarhat];
bbar=[NaN*ones(1,30) bbar];
bbarhat=[NaN*ones(1,30) bbarhat'];

figure(5)
subplot(2,2,1)                          % (2,2,2) for Panel C
plot(ybar,'b'), hold on
plot(ybarhat,'.b','LineWidth',2)
plot(abar,'g','LineWidth',1.5)
plot(abarhat,'om')
% plot(abarhat-bbarhat*(urefopt-10),'--k','LineWidth',1.5) % Don't use for Panel A
plot(ybarhatinit,'--b'), hold off, grid
ylabel('mean(y) and mean(a) [days10^-^1]')   % Don't use for Panel C
axis([30 60 -2 1])
text(30,1.2,'A','FontSize',14)          % C for Panel C

subplot(2,2,3)                          % (2,2,4) for Panel D
plot(bbar,'g','LineWidth',1.5), hold on
% plot(bbarhat,'.m','LineWidth',1.5)
plot(bbarhat,'om')
hold off, grid
xlabel('Generations')
ylabel('mean(b) [days10^-^1/^oC]')      % Don't use for Panel D
axis([30 60 -0.65 -0.45])
text(30,-0.435,'B','FontSize',14)       % D for Panel D

%% Results
opt_resultsresult=opt_resultsresult;
Mean=mean(opt_resultsresult)
Std=std(opt_resultsresult)
Mean_abar_error=mean(Rel_total_abar_error);
Std_abar_error=std(Rel_total_abar_error);
abar_results=[Mean_abar_error Std_abar_error]
Mean_bbar_error=mean(Rel_total_bbar_error);
Std_bbar_error=std(Rel_total_bbar_error);
bbar_results=[Mean_bbar_error Std_bbar_error]

Mean_abar_error_marked=mean(Rel_total_abar_error_marked);
Std_abar_error_marked=std(Rel_total_abar_error_marked);
abar_results_corr=[Mean_abar_error_marked Std_abar_error_marked]

Mean_fval=mean(Fvalresult);
Std_fval=std(Fvalresult);
fval_results=[Mean_fval Std_fval]

countW
countfval

%% Objective function
function f = fun_objective_Example(par,par_known,u,Y,ybar,W,Wbar,ybarhat_init,N,T)
Gbb=par(1);
Gab=par(2);
vare=par(3);
varw=par(4);
bbarhat_init=par(5);
uref=par(6);
Gaa=par_known(1);
ue=u-uref;
bbarhat=bbarhat_init*ones(T-31,1);
ybarhatobj=ybarhat_init*ones(T-31,1);

for t=1:29
    abarhat(1)=ybar(1)-bbarhat(1)*ue(1);
    Pyy=Gaa+vare+2*Gab*ue(t)+ue(t)^2*(Gbb+varw);
    Gyy=Gaa+Gab*ue(t)+Gab*ue(t+1)+ue(t)*Gbb*ue(t+1);
    covWy=(N-1)*cov(W(:,t),Y(:,t))/N;
    betay=inv(Pyy)*covWy(1,2)/Wbar(t);
    abarhat(t+1)=abarhat(t)+(Gaa+Gab*ue(t))*betay;
    bbarhat(t+1)=bbarhat(t)+(Gab+Gbb*ue(t))*betay;
    ybarhatobj(t+1)=abarhat(t+1)+bbarhat(t+1)*ue(t+1);
end
     f=sum((ybar(1:29)'-ybarhatobj(1:29)).^2);
end

%% Constraints function
function [cineq,ceq]=fun_constraints_Example(par,par_known,u1,u2,Y,ybar1,ybar2,W,Wbar,ybarhat_init,N,T)
cineq =  [];   % Compute nonlinear inequalities.
ceq = [];   % Compute nonlinear equalities.
end
```
